# Comparison of thigh muscle characteristics between older and young women using tensiomyography

**DOI:** 10.1101/2022.08.05.502971

**Authors:** Jung Hoon Chai, Chul-Hyun Kim, Sang-Won Bae

## Abstract

Tensiomyography is a non-invasive method of evaluating neuromuscular function through skeletal muscle contraction. The objective of this study was to compare the thigh muscle characteristics of older and young women using tensiomyography. Nineteen older and fifteen young women without musculoskeletal diseases were included. For the quadriceps, the bilateral vastus medialis (VM), vastus lateralis (VL), and rectus femoris (RF) and for the hamstrings, the bilateral semitendinosus (ST) and biceps femoris (BF) were measured. Result variables—maximal displacement (Dm), contraction time (Tc), and contraction velocity (Vc)—were compared. Dm values of the hamstrings of both legs and their summed values were significantly smaller in older women than in young women; no difference was found in the Dm values of the quadriceps. Tc and Vc of the hamstrings, VM, and VL were longer and slower, respectively, in older women than in young women. There were no significant differences in the Dm, Tc, or Vc of the RF between older and young women. Decreased Dm of the hamstrings in older women occurred due to changes in muscle function, but not muscle mass. The changes in the Tc of the hamstrings, VM, and VL indicate that type II muscle fibers were converted to type I in older women. There was no difference in RF between older and young women, implying that the RF is not affected by age. Our findings indicate that resistance exercises, which preserve the type II fibers, and flexibility exercises, which reduce stiffness, are appropriate for the lower extremity in older women.

## Introduction

During the aging process, the size and mitochondrial content of skeletal muscles decline while intramuscular adipose tissue accumulates, leading to muscular atrophy [1–3]. This reduction in muscular mass and strength due to aging is defined as sarcopenia [4]. Sarcopenia of the quadriceps, which often occurs to a serious degree in the older population [5], is a risk factor for falls and functional impairment [6,7]. This indicates that a decrease in muscular mass and strength due to aging can lead to a decline in muscular function.

Tensiomyography (TMG) was introduced as a device that evaluates muscles [8,9]. In particular, it is a non-invasive evaluation method that can evaluate muscular function as well as damage [10–12]. TMG induces muscle contraction through electrical stimulation and then measures the contraction time, which is the time it takes for the muscle belly to be maximally displaced. Maximal displacement (Dm), an output parameter obtained when performing TMG, has been reported to be highly correlated with muscle tone and stiffness [13,14]. Pisot et al. conducted a 35-day bed rest experiment and reported that the Dm of the biceps remained unchanged, whereas it was increased in the thigh muscles (gluteus maximus, vastus medialis [VM], and biceps femoris [BF]) [14]. This indicates that increases in Dm can potentially serve as an indicator of muscular atrophy.

Along with the Dm, contraction time (Tc) also constitutes a TMG output parameter; it is mostly used as an indicator of the effects of training. Završnik et al. compared the biomechanical characteristics of the vastus lateralis (VL) and rectus femoris (RF), particularly the running speed and Tc, and reported that the Tc of the RF had a stronger correlation with running speed than that of the VL [15]. Zubac and Šimunič reported that an improvement in jumping abilities was associated with a shorter Tc after an 8-week plyometric training program [16]. Calculating the contraction velocity (Vc), obtained by dividing the Dm by the Tc, has also been reported in the literature as a way to measure the speed of muscle contraction [17–20], and Loturco et al. reported that the Vc should be appropriate for the evaluation of functional adaptation [21].

Studies have also evaluated the reliability of TMG measurements [10,22], although the cut-off values for the normal ranges of the Dm and Tc, which are commonly reported in the literature, have not yet been established. Moreover, most studies have induced damage to subjects or applied exercise programs, which should be taken into consideration as mediator variables; thus, it is often difficult to interpret the significance of increases or decreases in the Dm or Tc.

Therefore, this study aimed to evaluate muscles in a relaxed state without any intervention, and the changes in muscles due to aging were evaluated using TMG in terms of the Dm, Tc, and Vc of the thigh muscles of older and young women.

## Materials and methods

### Subjects

This study was approved by the the Institutional Review Board on Human Subjects Research and Ethics Committee of Soonchunhyang University (approval number 1040875-201710-BM-044). Subjects without musculoskeletal diseases were recruited, resulting in a total of 19 older women (age: 71.2 ± 4.9 years, height: 155.7 ± 4.6 cm, weight: 63.7 ± 10.7 kg) and 15 young women (age: 20.9 ± 2.1 years, height: 161.7 ± 4.0 cm, weight 53.6 ± 5.3 kg) participating in the study. All subjects were asked whether they were willing to participate in the study and the purpose and background of the study were explained. Those who were willing to participate were requested to complete a written informed consent form.

### Tools

The TMG S1 model consists of a stimulator, which electrically stimulates muscles, a sensor, which transmits the muscular response to a computer, and a software program, which enables the researcher to confirm muscle responses (Fig 1).

**Fig 1.**
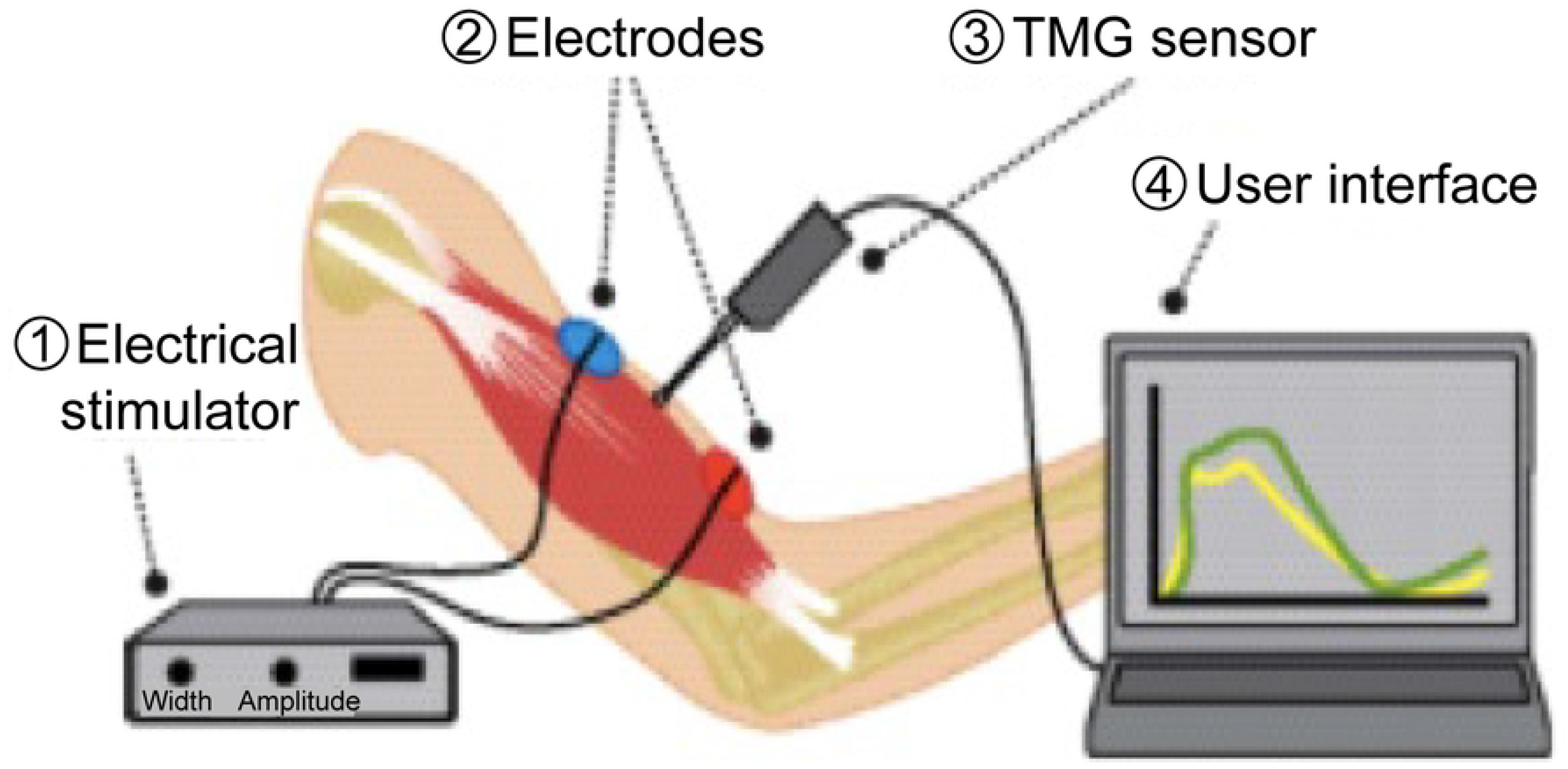
System components of TMG include. ① Electrical stimulator, ② electrodes, ③ TMG sensor, ④ User interface

Here, TMG presents the following five parameters: maximal displacement (Dm); delay time (Td) required to reach 10% of the Dm; contraction time (Tc) required to be between 10% and 90% of the Dm; sustain time (Ts) as the time between 50% of the muscle contraction and 50% of the relaxation; and relaxation time (Tr) as the time between 90% of the Dm and 50% after the peak of the Dm (Fig 2).

**Fig 2.**
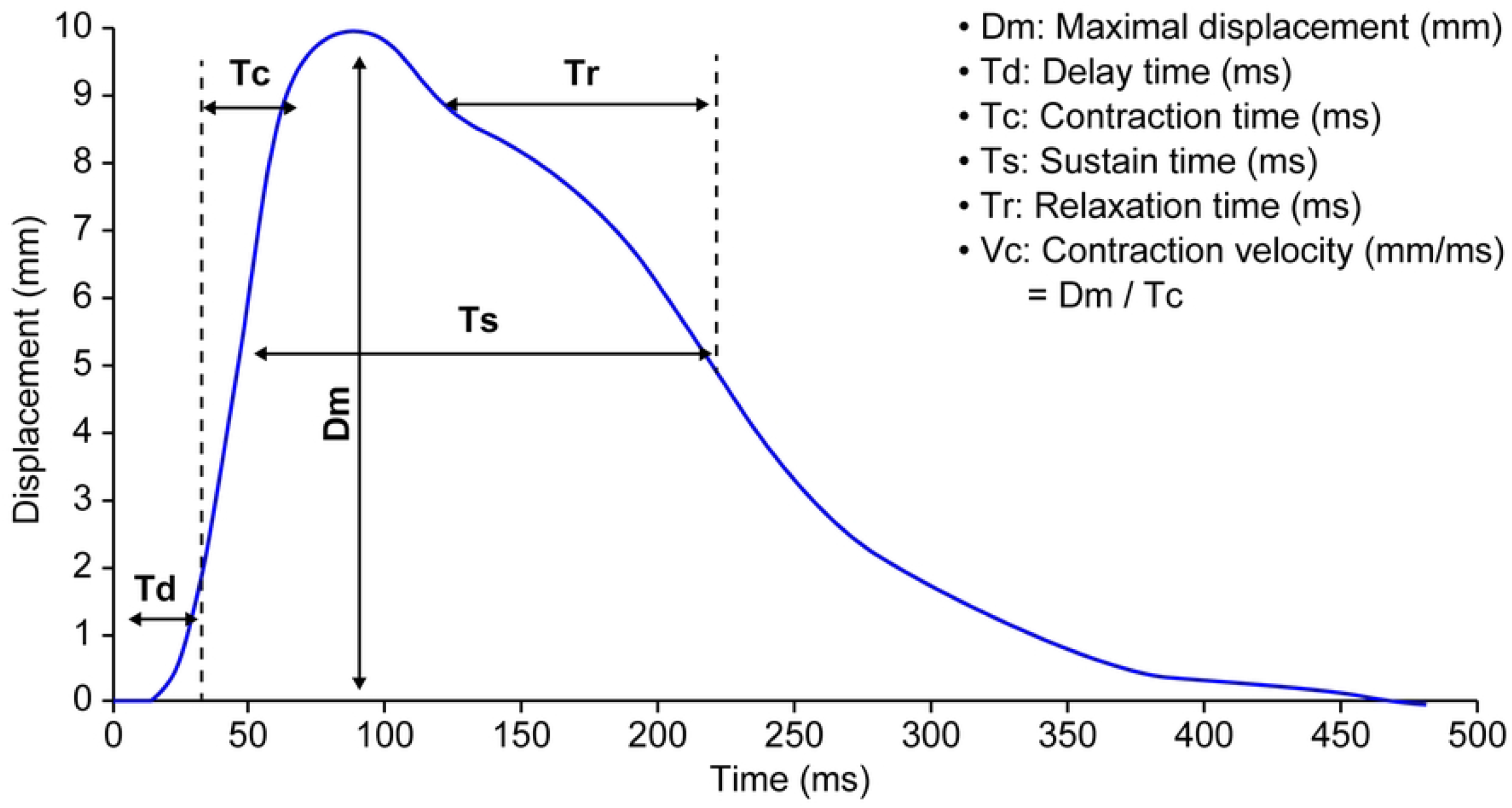
Tensiomyography and parameters.

### Measurement methods

The TMG sensor was placed on the belly of muscle to be measured, and an electrode was attached along the surface of the muscle 2–3 cm away from the sensor. The stimulation intensity was initially set at 20 mA and was increased in increments of 10 mA. The stimulation was continued until the maximum displacement of the muscle belly was observed, after which the test was continued in 10-second intervals to minimize potentiation and fatigue from previous stimulations.

The VM, VL, and RF of the quadriceps and the semitendinosus (ST) and BF of the hamstrings were selected for measurements, and the test was conducted bilaterally to account for dominance. The subjects laid in a supine position for measurement of the VM, VL, and BF, while maintaining their knees at 120 degrees; the ST and BF were measured in a prone position while maintaining the knees at 150 degrees. The measurements were compared in terms of the Dm, Tc, and Vc, where the Vc was obtained by dividing the Dm by the Tc.

### Data processing

Measurements were entered into Microsoft Excel (Microsoft, Redmond, WA, USA) and descriptive statistics were calculated (mean ± SD). Normality of the data was confirmed using SPSS for Mac Version 21.0 (IBM Co., Armonk, NY, USA), and independent t-tests were conducted to compare the data of the older and young women. A *p*-value < 0.05 was considered statistically significant.

## Results

The Dm, Tc, and Vc of the thigh muscles in young and older women are presented in Tables 1–3.

**Table 1.**
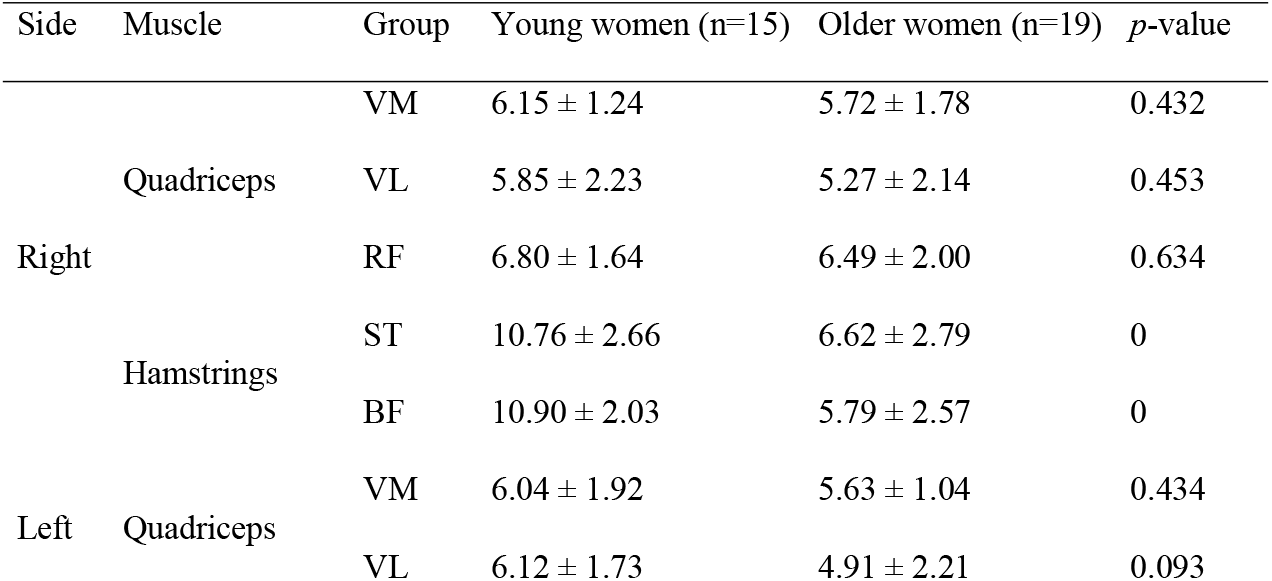
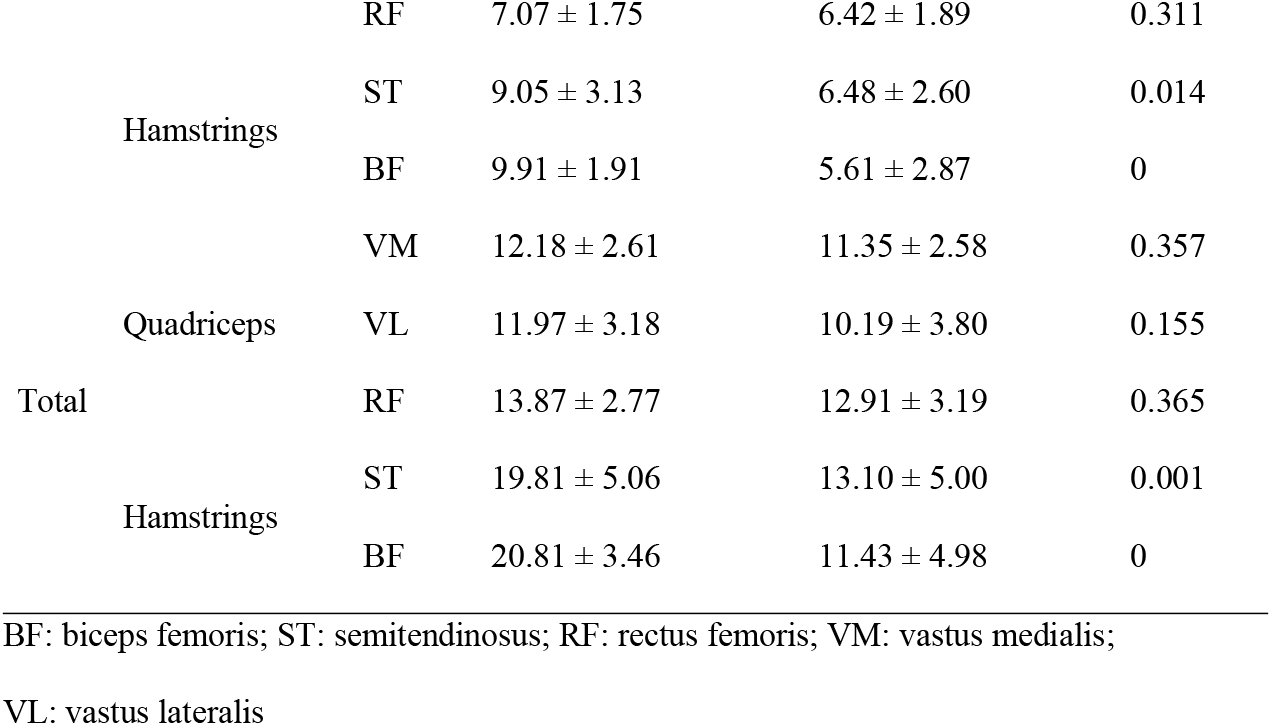
Maximal displacement (Dm) in both legs.

In the hamstrings, the Dm values obtained from the right and left sides, as well as that of the total, were significantly smaller in older women than in young women (p<0.05). In contrast, there was no difference in the Dm of the quadriceps between older and young women.

The Tc values of the hamstrings were significantly higher in the left, the right, and both legs in older women than in young women (p<0.05). The VM and VL of the quadriceps of older women had a significantly longer Tc in the left, the right, and both legs than had those of young women. However, there was no difference in the Tc of the RF between older and young women.

The left, right, and total Vc of the hamstrings were significantly slower in older women than in young women (p<0.05). The left, right, and total Vc of the VM and VL of the quadriceps were also slower in older women than in young women (p<0.05), and the right VL tended to be slower in older women than in young women (p=0.059). However, the Vc of the RF did not differ between older and young women (p>0.05).

## Discussion

This study aimed to quantitatively compare the function of the thigh muscles between older and young women using TMG, which can non-invasively measure muscle function. This is the first study to compare older women with young women in a TMG study without a specific intervention. Studies that used TMG have reported that Dm is a variable with a relatively small and stable change in value, and a decrease in Dm has been reported to be correlated with increases in muscle stiffness and tone [13,14,23,24]. Pisot et al. reported that Dm was increased in the atrophied thigh muscles by inducing muscular atrophy in healthy men on bed rest for 35 days, and increased Dm was associated with muscle atrophy and decreased muscle and tendon stiffness [14]. In addition, Chai et al. reported that the Dm in bodybuilder muscles was decreased compared to that in the general population and that this decrease was correlated with muscle hypertrophy [20]. In this study, the Dm of the hamstrings (ST, BF) decreased significantly in older women compared to that in young women, whereas no difference was observed in the quadriceps (VM, VL, RF) (Table 2).

**Table 2.**
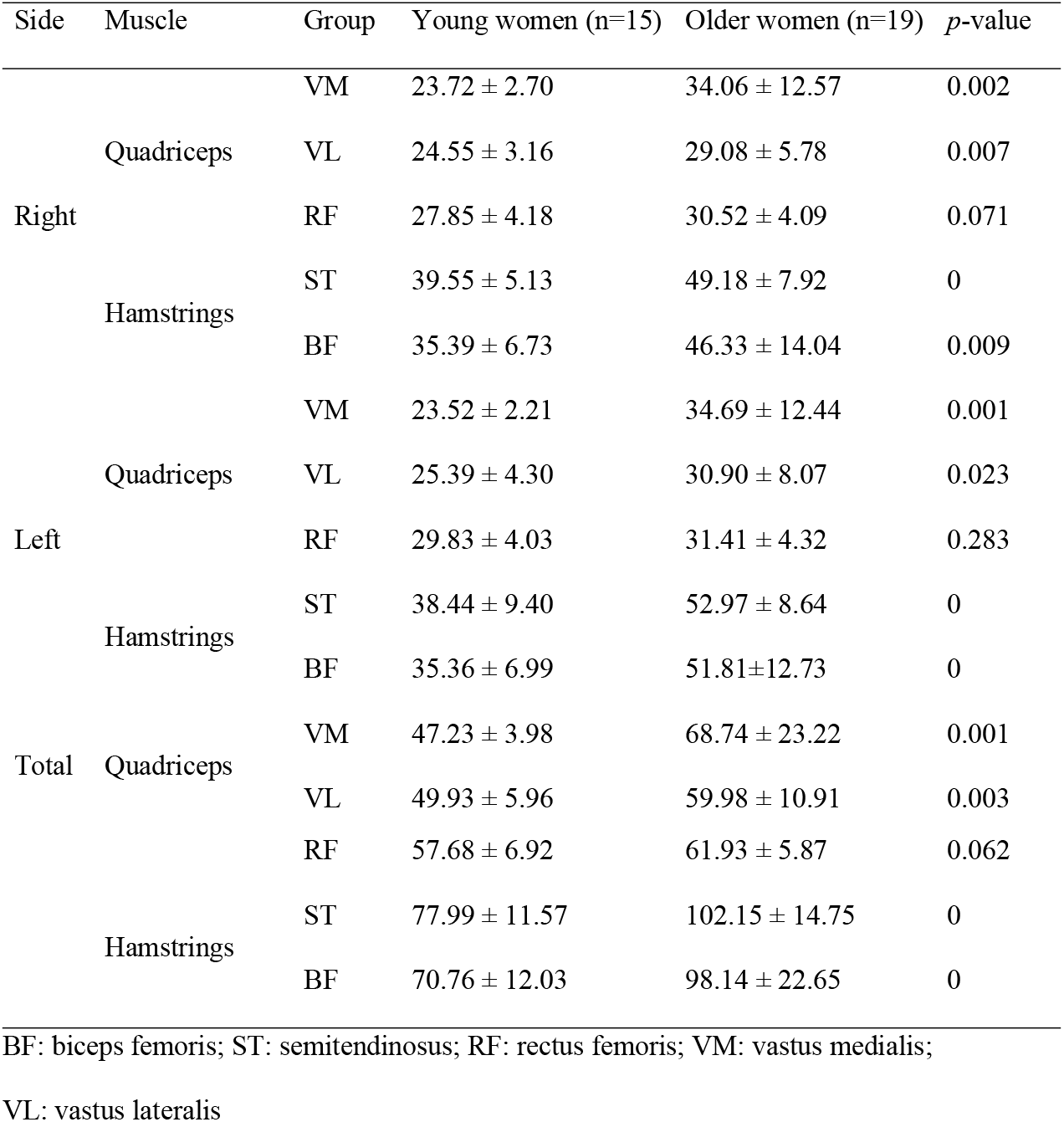
Contraction time (Tc) in both legs.

Aging is associated with a reduction in muscle and joint flexibility [25,26], which indicates increased stiffness [27,28]. As we age, the flexibility of our muscle and joints decreases [25,26] and stiffness increases [27,28].

A study that compared the femoral muscle mass and cross sectional area of older and young people using MRI and ultrasound reported no difference in the hamstrings between the groups, but it was reported that the quadriceps decreased in older individuals [5,29,30].

In this study, there was no significant difference in the Dm of the quadriceps between older and young women. This could be due to the offsetting effect of the increase in muscle stiffness (decrease in Dm) and the progression of muscle atrophy (increase in Dm) that occur in the quadriceps of older people. However, in the hamstring, the Dm decreased only by increase in stiffness without the effect of muscle atrophy.

In most of the thigh muscles, except the RF (VL, VM, ST, and BF), the Tc was higher in older women than in young women. The Tc, which is the time required for muscles to contract in response to an electrical stimulus, was reported to be highly correlated with the ratio (%) of type I muscle fibers [13]. It has also been used as a variable to evaluate the muscles of athletes as well as the effects of training [16,31]. The Tc of the thigh muscles was significantly longer in older women than in young women (except for that of the RF). A reduction in muscle mass due to aging can often be attributable to a decrease in type II fibers [32]. This indicates that the ratio of type I fibers to type II fibers should be higher in older people, which could explain an increased Tc, as seen in this study. Šimunič et al. used TMG to compare the VL, gastrocnemius medialis, and BF of endurance athletes, power athletes, and non-athletes of different ages and reported that the Tc increased with age, regardless of the form of exercise [33]; this is similar to the findings from our study.

Vc is the value obtained by dividing Dm by Tc, and it can be explained as the contraction velocity of muscle by electrical stimulation. It is also a variable suggested to correct for Tc affected by Dm [17,19,20]. In this study, Vc appeared more slowly in older women than in young women (most of the femoral muscles, except for RF). It can be predicted that the aged muscles, except for RF, may become relatively fatigued or have reduced muscle function.

In this study, the Dm decreased in the hamstrings of older women, whereas the Tc increased, resulting in substantial reduction of Vc (p<0.000). In contrast, in the quadriceps, the Dm of older women did not differ from that of young women, whereas the Tc increased. Thus, the Vc decreased in older women. Lockhart and Kim reported that older populations tended to have reduced hamstring activation [34]. In other words, the decrease in the Dm of the hamstrings that was observed in this study can be attributed to reduced hamstring activation in older people.

Another important finding of this study was regarding the features of the RF, which showed no difference between young and older women in terms of the Dm, Tc, or Vc (Tables 1–3). Miokovic et al. reported heterogeneous patterns in the lower extremity atrophy of particular muscles after 60 days of bed rest and reported that the RF underwent less atrophy than the vasti muscles [35]. In addition, Watanabe et al. suggested that the muscle thickness of RF in older women decreased compared to that in young people, but there was no difference in muscle activity according to electromyography [36]. This suggests that there is no change between type I and type II in RF, even with age. Through biopsy studies, aging causes changes in type II fibers in various muscles, but this has not been clearly explained in RF [37]. These previous findings as well as the results of the present study indicate that the RF maintains its function even with aging. Therefore, in future studies, it will be necessary to evaluate the change in form or structure in addition to the RF function.

**Table 3.**
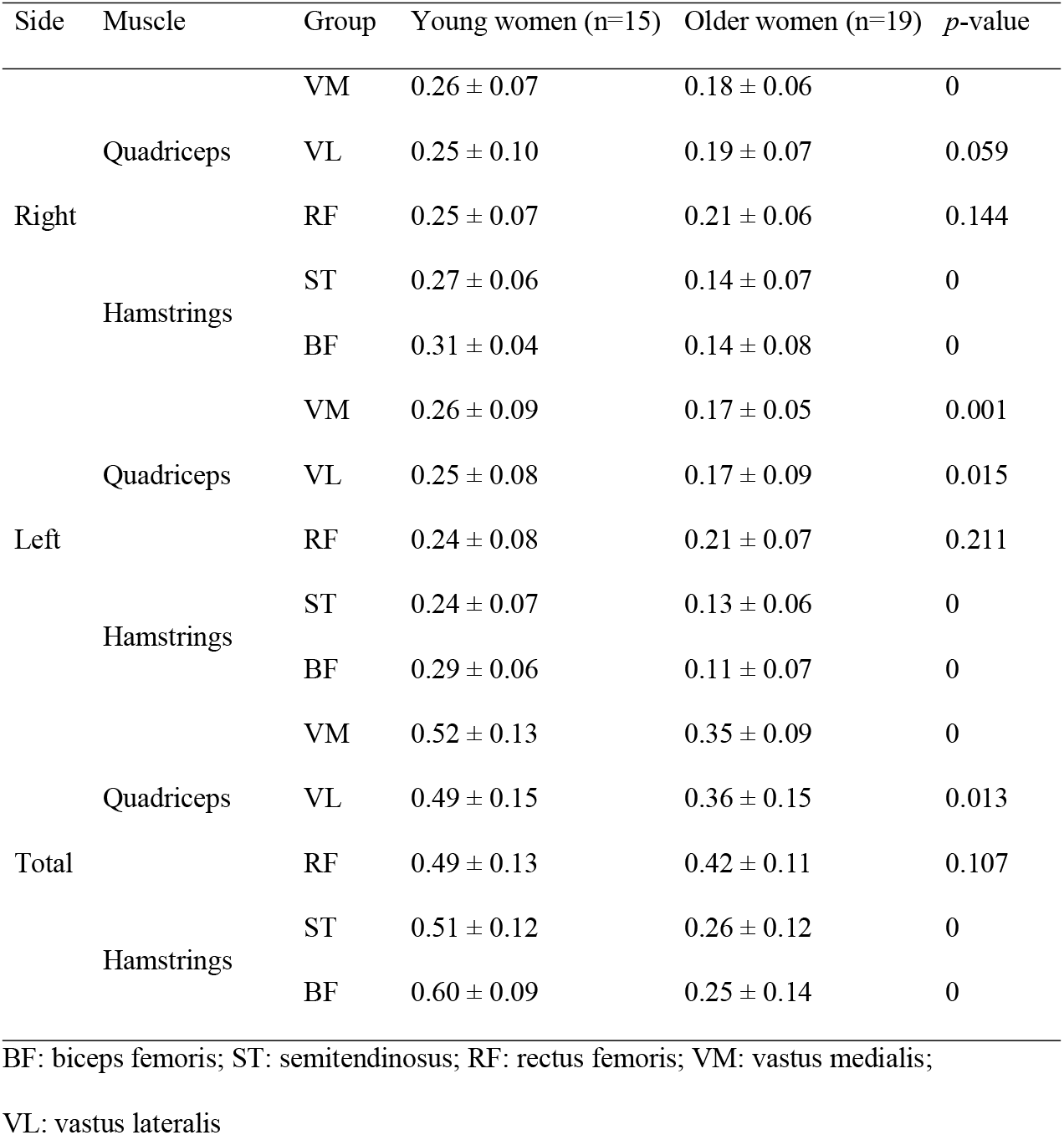
Contraction velocity (Vc) in both legs.

Recently, various methods, including strength evaluation through isokinetic testing and measurement of the cross-sectional area of the thigh muscles using computed tomography, have enabled the qualitative evaluation of muscles in the older population [38–40]. Although previous studies using TMG were mostly conducted among patients or athletes with muscle damage, this study showed that TMG could also evaluate muscle function in older people. In other words, TMG will enable the qualitative evaluation of muscles in older people.

In conclusion, this study confirmed, through TMG, that the hamstrings of older women are weaker than those of young women, whereas the RF function was relatively well preserved. These findings indicate that the hamstrings underwent a greater functional decline with aging (greater decrease in the Vc). The functional decline of the quadriceps was distinct from that of the hamstrings, which could have resulted from morphological decline owing to muscle atrophy. Meanwhile, the RF was found to have undergone a relatively less morphological and functional decline.

Finally, to prevent muscle atrophy and reduce the decrease and change of skeletal muscle type II fibers in older women, an exercise program consisting of resistance training, including stretching exercises, which are considered necessary to prevent stiffness, would be appropriate. In a future study, it will be necessary to develop and apply an exercise program that combines resistance and flexibility exercises suitable for older women to verify its effectiveness using TMG.

## Acknowledgements

None.

## References

1. Akima H, Yoshiko A, Hioki M, Kanehira N, Shimaoka K, Koike T, et al. Skeletal muscle size is a major predictor of intramuscular fat content regardless of age. Eur J Appl Physiol. 2015; 115:1627–1635.

2. Crane JD, Devries MC, Safdar A, Hamadeh MJ, Tarnopolsky MA. The effect of aging on human skeletal muscle mitochondrial and intramyocellular lipid ultrastructure. J Gerontol A Bio Sci Med Sci. 2010;65A: 119–128.

3. DeNino WF, Tchernof A, Dionne IJ, Toth MJ, Ades PA, Sites CK, et al. Contribution of abdominal adiposity to age-related differences in insulin sensitivity and plasma lipids in healthy nonobese women. Diabetes Care. 2001;24: 925–932.

4. Rosenberg IH. Sarcopenia: origins and clinical relevance. J Nutr. 1997;127: 990s–9901s.

5. Ogawa M, Yasuda T, Abe T. Component characteristics of thigh muscle volume in young and older healthy men. Clin Physiol Funct Imaging. 2012;32: 89–93.

6. Pijnappels M, van der Burg PJ, Reeves ND, van Dieën JH. Identification of elderly fallers by muscle strength measures. Eur J Appl Physiol. 2008;102: 585–592.

7. Van Roie E, Verschueren SM, Boonen S, Bogaerts A, Kennis E, Coudyzer W, et al. Force-velocity characteristics of the knee extensors: an indication of the risk for physical frailty in elderly women. Arch Phys Med Rehabil. 2011;92: 1827–1832.

8. Macgregor LJ, Hunter AM, Orizio C, Fairweather MM, Ditroilo M. Assessment of skeletal muscle contractile properties by radial displacement: the case for tensiomyography. Sports Med. 2018;48: 1607–1620.

9. Martín-Rodríguez S, Alentorn-Geli E, Tous-Fajardo J, Samuelsson K, Marín M, Álvarez-Díaz P, et al. Is tensiomyography a useful assessment tool in sports medicine? Knee Surg Sports Traumatol Arthrosc. 2017;25: 3980–3981.

10. Wilson HV, Johnson MI, Francis P. Repeated stimulation, inter-stimulus interval and inter-electrode distance alters muscle contractile properties as measured by tensiomyography. PLoS One. 2018;13: e0191965.

11. Kim C, Chai JH, Kim BK, Kim CH, Bae SW. A novel method for the assessment of muscle injuries. Korean J Sports Med. 2015;33: 59–66.

12. Chai JH, Kim C, Kim CH. TMG (tensiomyography): non-invasive method of evaluation of muscle function. Korean J Phys Educ. 2017;56: 519–526.

13. Dahmane R, Valenčič V, Knez N, Eržen I. Evaluation of the ability to make non-invasive estimation of muscle contractile properties on the basis of the muscle belly response. Med Biol Eng Comput. 2001;39: 51–5.

14. Pisot R, Narici MV, Simunic B, De Boer M, Seynnes O, Jurdana M, et al. Whole muscle contractile parameters and thickness loss during 35-day bed rest. Eur J Appl Physiol. 2008;104: 409–414.

15. Završnik J, Pišot R, Šimunič B, Kokol P, Blažun Vošner H. Biomechanical characteristics of skeletal muscles and associations between running speed and contraction time in 8-to 13-year-old children. J Int Med Res. 2017;45: 231–245.

16. Zubac D, Šimunič B. Skeletal muscle contraction time and tone decrease after 8 weeks of plyometric training. J Strength Cond Res. 2017;31: 1610–1619.

17. García-García O, Serrano-Gómez V, Hernández-Mendo A, Morales-Sánchez V. Baseline mechanical and neuromuscular profile of knee extensor and flexor muscles in professional soccer players at the start of the pre-season. J Hum Kinet. 2017;58: 23–34.

18. García-Manso JM, Rodríguez-Matoso D, Sarmiento S, de Saa Y, Vaamonde D, Rodríguez-Ruiz D, et al. Effect of high-load and high-volume resistance exercise on the tensiomyographic twitch response of biceps brachii. J Electromyogr Kinesiol. 2012;22: 612–619.

19. Rodríguez-Ruiz D, Diez-Vega I, Rodríguez-Matoso D, Fernandez-del-Valle M, Sagastume R, Molina JJ. Analysis of the response speed of musculature of the knee in professional male and female volleyball players. Biomed Res Int. 2014;2014: 239708.

20. Chai JH, Kim BK, Kim C, Kim CH, Bae SW. Analysis of bodybuilder’s skeletal muscle characteristics using tensiomyography. Korean J Sports Med. 2016;34: 146–152.

21. Loturco I, Pereira LA, Kobal R, Kitamura K, Ramírez-Campillo R, Zanetti V, et al. Muscle contraction velocity: a suitable approach to analyze the functional adaptations in elite soccer players. J Sports Sci Med. 2016;15: 483–491.

22. Martín-Rodríguez S, Loturco I, Hunter AM, Rodríguez-Ruiz D, Munguia-Izquierdo D. Reliability and measurement error of tensiomyography to assess mechanical muscle function: a systematic review. J Strength Cond Res. 2017;31: 3524–3536.

23. Valenčič V, Knez N. Measuring of skeletal muscles’ dynamic properties. Artif Organs. 1997;21: 240–242.

24. Rey E, Lago-Peñas C, Lago-Ballesteros J. Tensiomyography of selected lower-limb muscles in professional soccer players. J Electromyogr Kinesiol. 2012;22: 866–872.

25. Golding LA, Lindsay A. Flexibility and age. Perspective. 1989;15: 28–30.

26. Mazzeo R, Cavanagh P, Evans WJ, Fiatarone M, Hagberg J, McAuley E, Startzell J. ACSM position stand: exercise and physical activity for older adults. Med Sci Sports Exerc. 1998;30: 992–1008.

27. Hortobágyi T, DeVita P. Altered movement strategy increases lower extremity stiffness during stepping down in the aged. J Gerontol A Biol Sci Med Sci. 1999;54: B63–B70.

28. Hsu MJ, Wei SH, Yu YH, Chang YJ. Leg stiffness and electromyography of knee extensors/flexors: comparison between older and younger adults during stair descent. J Rehabil Res Dev. 2007;44: 429–435.

29. Young A, Stokes M, Crowe M. Size and strength of the quadriceps muscles of old and young women. Eur J Clin Invest. 1984;14: 282–287.

30. Fukumoto Y, Ikezoe T, Yamada Y, Tsukagoshi R, Nakamura M, Takagi Y, et al. Age-related ultrasound changes in muscle quantity and quality in women. Ultrasound Med Biol. 2015;41: 3013–3017.

31. Rusu LD, Cosma GG, Cernaianu SM, Marin MN, Rusu PF, Ciocănescu DP, et al. Tensiomyography method used for neuromuscular assessment of muscle training. J Neuroeng Rehabil. 2013;10: 67.

32. Nilwik R, Snijders T, Leenders M, Groen BB, van Kranenburg J, Verdijk LB, et al. The decline in skeletal muscle mass with aging is mainly attributed to a reduction in type II muscle fiber size. Exp Gerontol. 2013;48: 492–498.

33. Šimunič B, Pišot R, Rittweger J, Degens H. Age-related slowing of contractile properties differs between power, endurance, and nonathletes: a tensiomyographic assessment. J Gerontol A Biol Sci Med Sci. 2018;73: 1602–1608.

34. Lockhart TE, Kim S. Relationship between hamstring activation rate and heel contact velocity: factors influencing age-related slip-induced falls. Gait Posture. 2006;24: 23–34.

35. Miokovic T, Armbrecht G, Felsenberg D, Belavý DL. Heterogeneous atrophy occurs within individual lower limb muscles during 60 days of bed rest. J Appl Physiol. 2012;113: 1545–1559.

36. Watanabe K, Kouzaki, M, Moritani, T. Effect of aging on region-specific functional role and muscle geometry along human rectus femoris muscle. Muscle Nerve. 2017;56: 982–986.

37. Brunner F, Schmid A, Sheikhzadeh A, Nordin M, Yoon J, Frankel V. Effects of aging on type II muscle fibers: A systematic review of the literature. J Aging Phys Act. 2007;15: 336–348.

38. Fragala MS, Kenny AM, Kuchel GA. Muscle quality in aging: a multi-dimensional approach to muscle functioning with applications for treatment. Sports Med. 2015;45: 641–658.

39. Moore AZ, Caturegli G, Metter EJ, Makrogiannis S, Resnick SM, Harris TB, et al. Difference in muscle quality over the adult life span and biological correlates in the Baltimore Longitudinal Study of Aging. J Am Geriatr Soc. 2014;62: 230–236.

40. Yoshiko A, Hioki M, Kanehira N, Shimaoka K, Koike T, Sakakibara H, et al. Three-dimensional comparison of intramuscular fat content between young and old adults. BMC Med Imaging. 2017;17: 12.

